# A precise and general FRET-based method for monitoring structural transitions in protein self-organization

**DOI:** 10.1101/2021.02.25.432866

**Authors:** Qi Wan, Sara N. Mouton, Liesbeth M. Veenhoff, Arnold J. Boersma

## Abstract

Proteins assemble into a tremendous variety of dynamic and functional structures. Sensitive measurements directly in cells with a high spatiotemporal resolution are needed to distinguish these different assemblies. Here, we demonstrate precise and continuous monitoring of cytoplasmic protein self-assemblies and their structural transitions. Intermolecular FRET with both the donor and acceptor protein at the same target protein provides high sensitivity while retaining the advantage of straightforward ratiometric imaging. We measure different assembly structures, transient intermediate states’ kinetics, and assembly formation resolved in space and time. Specifically, the method recapitulates that i) the mutant Huntingtin exon1 (mHttex1) protein first forms low-FRET and presumably less ordered assemblies in yeast and human cells, which develop into high-FRET aggregates, ii) the chaperone DNAJB6b prevents low-FRET mHttex1 assemblies, yet coassembles with mHttex1 aggregates, and iii) FUS’ condensates have mutation-dependent nanoscopic structures. FACS measurements allow assembly measurement in a high-throughput manner crucial for screening efforts, while fluorescence microscopy provides spatiotemporally-resolved measurements on the single-condensate level during a cell’s lifetime to assess the biological consequences. Implementation in other native or non-native proteins could provide insight into many studies involving protein condensation or aggregation.

## Introduction

Proteins assemble into different structures that range from quaternary structures such as the cytoskeleton and multi-subunit enzymes to dynamic protein condensates and misfolding-induced aggregates. The structure and dynamics of condensates and aggregates depend on the local biochemistry. For example, aggregate-forming proteins associated with, e.g., Parkinson’s, Alzheimer’s, and Huntington’s disease have structures and aggregation pathways resulting from chaperone activity, posttranslational modifications, the inclusion of other biomolecules, and the cell’s physicochemical properties.^1-4^ Thus, aggregate structures vary considerably, and better monitoring of the structures may improve identifying the best drug targets. Condensate structures also vary: The protein Fused in Sarcoma (FUS) forms dynamic liquid-like assemblies, but also aberrant gel-like states mislocalized in the cytoplasm sometimes associated with Amyotrophic lateral sclerosis.^5,6^ To better understand these condensates’ biological function and related diseases, it is essential to obtain insight into the structural transitions that protein assemblies undergo.

The structural transitions of protein assemblies are challenging to determine inside cells. In buffer, reconstitution of purified protein allows detailed study of condensates and aggregates, but results may not be relevant in cells. Chemical or cryofixation of cells allows antibody staining and electron microscopy, providing the aggregate’s microscopic structure and its location in the cell. However, to understand what structures are present in living cells, how fast they form, their dynamics, and whether transient intermediates form, measurements inside living cells are required. The dynamics of protein assembly can be observed by tagging the corresponding proteins with a fluorescent protein or by the addition of small-molecule fluorescent dyes such as Thioflavin T,^7^ which usually only indicate amyloid-type structures. The target protein fused with a fluorescent protein allows fluorescent microscopy with a high spatiotemporal resolution, observation of dynamic behavior, and colocalization with other fluorescently tagged molecules.

Förster Resonance Energy Transfer is an ideal method to probe events at the 2-10 nm scale, which is in a protein’s size range. The FRET efficiency between a donor and acceptor fluorescent protein increases with an inverse distance to the power six when the fluorophores are moving closer towards each other. The orientation between the fluorophores alters the FRET efficiency additionally. Thus, FRET has found extensive use in determining protein-protein interactions and observing protein aggregation, such as the mutant Huntingtin exon 1 fragment containing an extended polyglutamine domain (mHttex1).^8-10^ To this end, a fluorescent donor is fused to mHttex1 and the acceptor to the second copy of mHttex1. Cotransfection of both plasmids allows the observation of an increase in FRET efficiency as both copies coaggregate with a short intermolecular distance in between. Unequal expression of donor and acceptor proteins can be corrected for using correction factors.^11^ Alternative methods have been developed recently, such as partial photo converting a GFP into an RFP to generate FRET^12^ or determine the aggregate’s influence on GFP’s fluorescent properties.^13-15^ However, the equal number of donors and acceptors in every cell is not achieved, or quantitative ratiometric imaging is not possible. These drawbacks can be overcome by fusing both the donor and acceptor to the same protein. However, fusing mHttex1 between a FRET pair impedes mHttex1 aggregation and depends on both inter and intramolecular FRET changes.^16,17^ Hence, although FRET is a beneficial tool to observe protein assembly, various hurdles remained in vivo to reach its full potential in cells.

Here we demonstrate straightforward and precise monitoring of condensate and aggregate properties with a FRET-based method that relies on conjugating the target protein to a fusion that contains both the donor and acceptor. We validate the method’s application by observing different states in single mHttex1 aggregates, the role and the oligomeric state of DNAJB6 during mHttex1 aggregation, and the differences in nanoscopic condensate structures formed by FUS mutants, all with the high spatiotemporal resolution that confocal fluorescence microscopy allows.

## Results

### Sensor design and approach

We aimed to develop a platform for efficient and precise intermolecular FRET measurements by fusing both the donor and acceptor to the same protein. To prevent intramolecular FRET changes during assembly, we adopted a FRET pair previously used for monitoring protein clustering in cell membranes.^18^ Ma et al. removed the native linker domains of the mVenus and mCherry proteins (VC) so that the FRET pair adopts a rigid conformation and stabilizes intramolecular FRET to a low level. Hence we argued that fusing this pair to condensate and aggregate forming proteins would reveal insights into short-lived intermediate states and their kinetics by following the intermolecular FRET efficiency in time. Intermolecular FRET increases with protein concentration, which can be measured by direct acceptor (mCherry) excitation. We expect this dependence to be characteristic for a specific condensate’s structural properties: While freely moving fluorophores interact randomly, a preferred binding-site provides a dominant orientation and distance between the fluorophores. For example, at the same protein concentration, a binding mode where the fluorophores are close together would provide a higher FRET than randomly mixed proteins. Hence this approach will be sensitive to changes in the relative distance between the VC domains and their respective orientation. Thus, the method allows distinction between different ordered states based on the FRET efficiency.

### mHttex1 self-assembly in human cells

We fused the VC domain first to the well-studied mutant Huntingtin Exon 1 fragment (mHttex1) containing an extended polyglutamine stretch of Q71, the native N-terminal peptide, and the native proline-rich domain to generate Q71-VC (Fig. 1A).^19^ We fused the VC at the C-terminus of mHttex1 because the fusion of mHttex1 with GFP has been characterized extensively. As observed by confocal fluorescence microscopy, transfection and expression of Q71-VC in HEK293t cells lead to the corresponding aggregates (Fig. 1B). Subsequent cell lysis and SDS-PA gel electrophoresis showed that the constructs were expressed intact and generated the typical SDS-insoluble aggregates (Supp. Fig. 1C). To confirm the FRET mechanism and the expected aggregate’s static properties, we determined the fluorescence recovery after photobleaching (FRAP) the acceptor (Supp. Fig. 1A,B). Indeed, bleaching the acceptor gave a substantial increase in donor emission, confirming high FRET efficiency. Moreover, recovery of the fluorescence did not occur as expected from mHttex1 aggregates. Hence, the fusion of VC to mHttex1 does not impede its typical aggregation behavior and gives high FRET efficiency.

**Figure 1:**
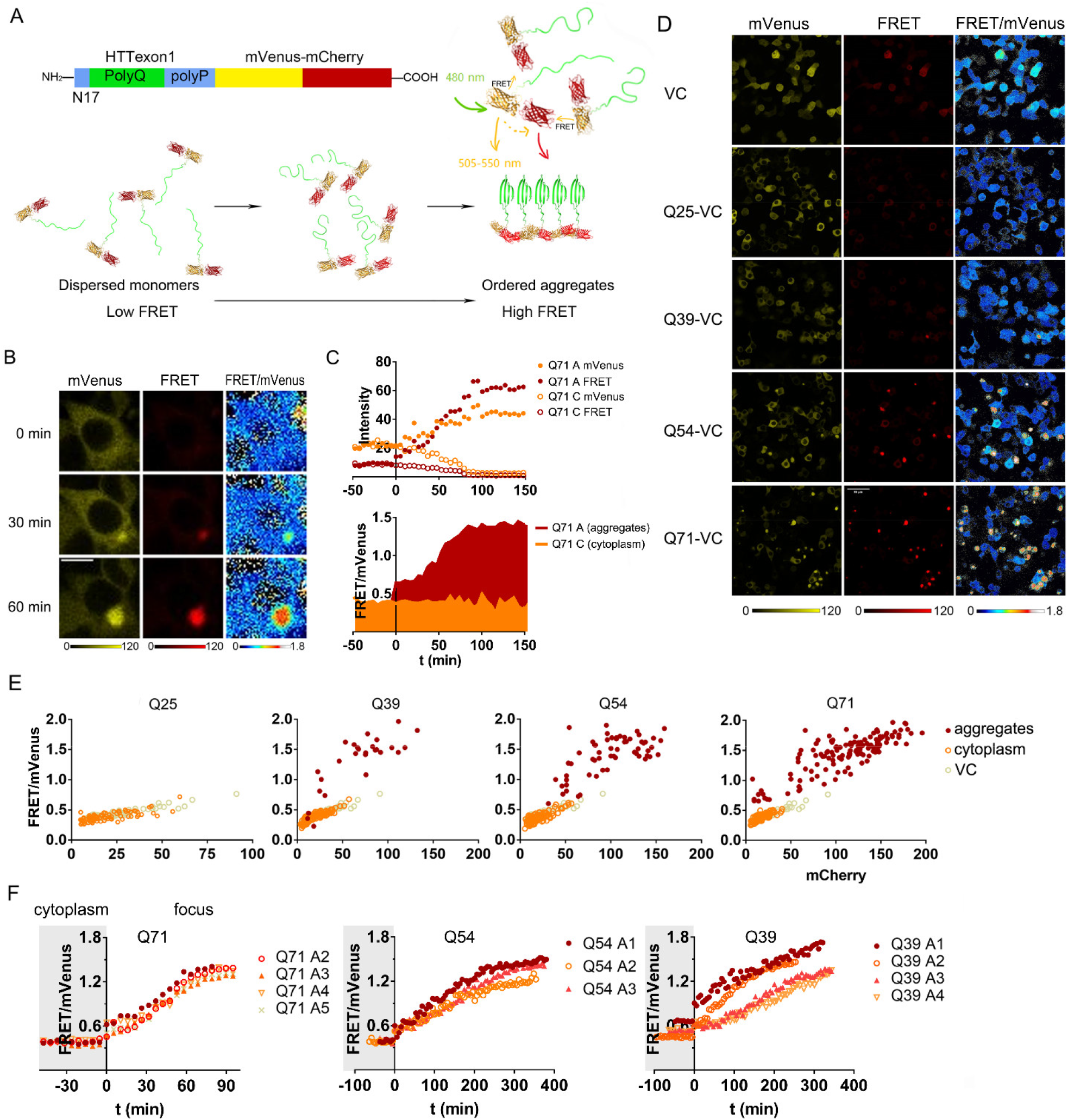
Tagging mHttex1 with the VC domain allows detailed observation of the aggregation progression in HEK293t. A. Design of the probe and cartoon of the sensing mechanism. The mVenus-mCherry FRET pair (VC) is fused to the C-terminus of mHttex1. Closer association between mHttex1 peptides increases the intermolecular FRET between the VC tags (N17: native N-terminal peptide, polyQ: polyglutamine, polyP: proline-rich domain). B. Fluorescence intensities and ratiometric images of Q71-VC in time measured by confocal microscopy. Excitation: 488 nm; emission: 505-555 nm, 600-700 nm. C. Quantifications of single-cell Q71-VC intensity in foci (A) and cytoplasm (C) (above) and the corresponding FRET ratios (below) over time. This cell displays a 20-min stable low FRET state. D. Fluorescence and FRET/donor ratiometric images of mHttex1 with different polyglutamine lengths. E. FRET/mVenus ratios of Q71, Q54, Q39, and Q25 labeled with VC plotted versus the mCherry intensity. The data for the foci, the cytoplasm, and the VC control are compared. Each data point is from a different cell. F. Progression of single-foci FRET ratios of Q71, Q54, Q39, and Q25-VC in time. The moment foci emerged was set as t=0; the data before t=0 is the FRET/mVenus ratio in the cytoplasm. Scale bars are 10 μm. A1-A5 are individual foci in different cells.

We quantified the readout by dividing the FRET channel fluorescence (excitation at 488 nm, emission 600-700 nm) by the donor channel (excitation at 488 nm, emission 505-555 nm) (Fig. 1C). Before aggregation, the FRET/donor ratio of Q71-VC is ∼0.4, equaling the control VC without Q71, while the Q71-VC FRET/donor ratio increases to 1.3-1.4 upon aggregation. The FRET/donor ratios are much less noisy than the donor emission alone and even provide a larger amplitude, therefore providing a higher sensitivity than simple fluorescent protein intensity measurements. The Q71-VC’s FRET/donor ratio remains low outside the aggregate and overlays with the VC control (Fig. 1E), with a similar small dependence of the FRET/donor ratio on the probe concentration due to intermolecular FRET. These data show that when Q71-VC is not in aggregates, it behaves as the VC, which contains monomeric variants mVenus and mCherry; hence Q71-VC is indistinguishable from a monomeric state when dispersed.

We analyzed the aggregation dependence on the number of glutamines by replacing the 71 glutamines of Q71-VC with 25 (Q25-VC), 39 (Q39-VC), and 54 (Q54-VC) glutamines. Longer glutamine stretches are known to increase aggregate formation, while the native 25 glutamine stretch does not aggregate inside cells. Indeed, we do not observe aggregates for Q25-VC under our conditions (Fig. 1D), which has FRET/donor ratios with an equal dependence on the concentration as the control VC (Fig. 1E). The Q39, Q54, and Q71 form foci with final ratios in the order of 1.3-1.4, suggesting that the Q length has little effect on the aggregate’s final structure in the 2-10 nm domain. Our method does not probe large morphological changes that could occur instead. The number of aggregates depends on the Q length: Q39-VC forms aggregates in only 5% of the cells and Q71-VC in 48%. We find that the onset of foci formation for the Q54-VC and Q71-VC occurs consistently at a similar concentration corresponding to 14±1 and 13±2 a.u. (s.d., n=4) of acceptor fluorescence, respectively. In contrast, the Q39-VC requires a ∼4 times higher concentration, i.e., 50±22 a.u. (s.d., n=4), with a higher variability between cells. Hence, the eventual aggregates have a similar structure, but longer Q stretches are more prone to aggregate.

The method allows for obtaining detailed kinetic information at the single-cell level (Fig. 1F). In most cases, the foci’ FRET/donor increase has a smooth and somewhat sigmoidal appearance; the aggregate’s structure evolves continuously without a strongly stabilized intermediate (low-FRET) state. In a single Q71-VC (shown in Fig. 1C) and two Q39-VC foci, however, the FRET/donor ratio appears to stabilize at a low level for up to 20-60 minutes before increasing. Fitting the data from the subsequent steep increase to an exponential decay allows comparing the aggregation kinetics and determine the apparent half-life t1/2 to aggregation. We find that Q71-VC transits to the final aggregates an order of magnitude faster than Q54-VC (t1/2 = 32±6 min (s.d.,n=5) and t1/2 = 244±77 min (s.d.,n=3), respectively). The Q71 aggregates both mature fastest and have a higher chance to form foci. Although the aggregates formed by Q39 appear to aggregate with similar kinetics as Q54 (t1/2 = 277±143 min (s.d.,n=3)), the concentrations of Q39 are ∼4× higher at the onset of foci formation, which accelerates aggregation. Together, the probes’ application in human cells allows observing single-cell aggregation kinetics and comparison between different Q-lengths and expression levels.

### Spatiotemporally resolved mHttex1 transition in yeast

The VC probing method can directly compare mHttex1 aggregation intermediates and kinetics between yeast and human cells: Yeast is a model organism to study mHttex1, albeit that yeast aggregates form solid inclusions instead of macroscopic fibrils.^20^ To this end, we loaded yeast in a microfluidics chip to follow aggregation under controlled environmental conditions (temperature, nutrients, pH) in individual cells.^21^ We transformed yeast with the plasmids encoding for the Q72-VC and the control VC and induced expression with 0.2% galactose on-chip. To resolve the different aggregation states, we measured individual cells every 20 minutes for 15 hours by widefield fluorescence microscopy (Fig. 2A,E). The control VC displayed ratios stable throughout the experiments with low dependence on the concentration (Supp. Fig. 2). For Q72-VC, on the other hand, the first foci appeared after 338±131 min (± s.d.). The size of the foci increased over time as expression continued and resulted in large aggregates or inclusions.

**Figure 2.**
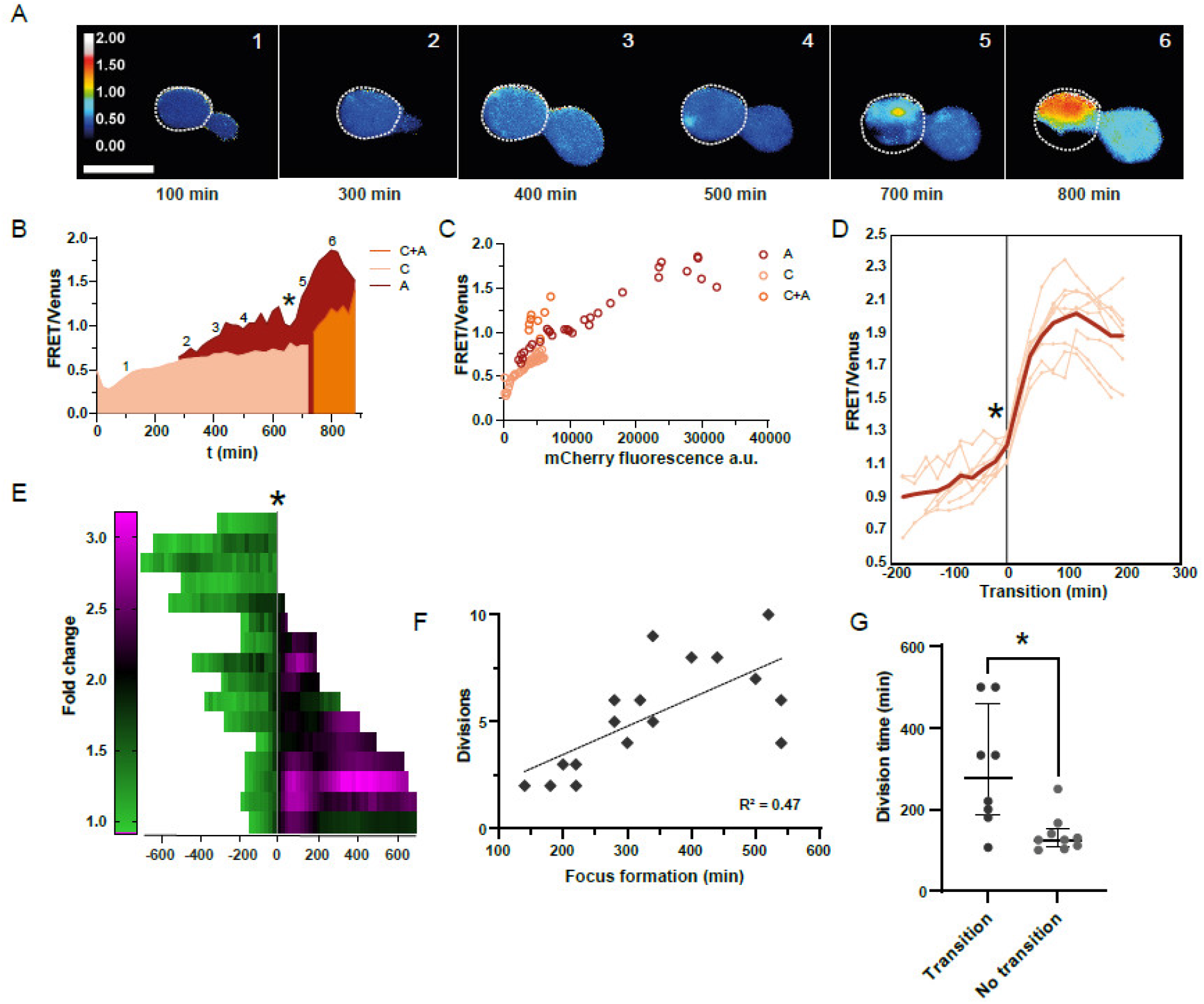
Long-term imaging of individual yeast cells reveals distinct phases of mHttex1 aggregate progression. A. Ratiometric images of a representative yeast cell expressing 72Q-VC. The cell outline is indicated with a white dotted line. The early low-FRET assemblies are visible in the Venus channel (not shown) but invisible in the ratiometric image as their FRET-ratio is similar to the cytosol. The white numbers correspond to the markings in panel B. Scale bar is 5 μm. B. FRET/mVenus ratios plotted against time for the cell displayed in A. The cytoplasm (C) is shown in pink and the aggregate (A) dark red. When additional small foci appear at later stages, a cytoplasmic region without aggregates is difficult to distinguish, and FRET/mVenus ratios are shown in orange (C+A). When the FRET/mVenus ratio in the aggregate starts to increase steeply is indicated with a star. C. FRET/mVenus ratios from the cytoplasmic, aggregate, and mixed regions plotted versus the mCherry signal. The acceptor’s fluorescence intensity is a direct indication of the probe concentration. D. Eight single-cell trajectories aligned to the timepoint (*) at which the FRET/mVenus ratios transition to higher FRET/donor values. The transition point was set as t=0, and trajectories ±200 min were plotted. The bold red line represents the average. E. Fold-change in aggregate FRET ratios plotted in time for each cell and aligned at the transition point indicated by a grey line and a star. F. The timing of the first appearance of the focus correlates with the division time plotted as the number of divisions completed by each cell during the experiment (R^2^ = 0.4713, p = 0.0023). G. Cells that harbor foci with lower FRET divide faster (Mann-Whitney test, p = 0.0101) than those transitioned to higher FRET foci.

When analyzing the single-cell traces in time, we find that the FRET/mVenus ratios in the assemblies first increase slowly, followed by a faster increase (Fig. 2B, Supp. Fig. 4). The star in Fig. 2B,E indicates the transition from the phase of slower increase to the phase of faster increase; the transition occurs at a FRET/mVenus ratio between 1.1 and 1.3. At FRET/mVenus ratios below 1.3, the ratio is concentration-dependent as assessed from direct acceptor excitation (Fig. 2C), while the FRET/mVenus becomes concentration-independent at the higher ratios after the transition. We interpret these changes in FRET/mVenus ratios as showing a structural transition between a more disorganized (low-FRET) early and a more organized (high FRET) late state of the Q-72-VC assembly. The length of the low-FRET state varies significantly with lifetimes between 100 and 700 min. When averaging a subset of the cells that display the transition to a high FRET/donor ratio (8 out of 16) (Fig. 2D, Supp. Fig. 3A), we observe that the aggregation proceeds through the typical aggregation curve with t1/2 = 42±4 min (± s.e., n=8). Thus, while cells vary in the low-FRET state’s length, they transit to the high-FRET state with similar kinetics once they enter the transition point. The remainder of the cells displays assemblies that do not transform into high-FRET assemblies (FRET/donor remains <1.5; Supp. Fig. 3B) during the duration of the experiment (15h); they may well do so at later timepoints. In contrast to mammalian cells, the FRET/donor ratio in the yeast high-FRET assemblies is unstable, which may result from additional structural rearrangements in the aggregates.

Assessing the ratiometric images (Fig. 2A), we see spatial heterogeneity in the FRET/donor ratios inside some of the aggregates along the aggregation trajectory, where assemblies with low FRET/donor already contain an area with the aggregate’s high FRET/donor (e.g., see Fig 2A, 700 minutes). Hence, the transition is temporally and spatially resolved and spreads throughout the assembly as expected after a nucleation event. Further, the ratios outside high FRET/donor aggregates also increase steadily (Fig. 2B), suggesting more widespread aggregation. Indeed, as aggregation proceeds, multiple visible aggregates appear in the cytoplasm, especially for cells with very high expression levels, as noted previously for yeast.^20^

The division time of a yeast cell is a readout of cell fitness. There is a natural variation in division times of genetically identical yeast cells grown under these well-controlled conditions.^22^ Interestingly, we find that the focus’s appearance’s timing correlates with the number of divisions a cell completes during the recording, where foci appear later in the faster-dividing cells (Fig. 2F). Moreover, we find that foci in faster dividing cells often remain at the low-FRET level (Fig. 2G). These correlations suggest that fitter cells have a higher protein homeostasis capacity which delays the appearance of the focus as well as the transition to the high FRET-state.

Together, the mHttex1 aggregation monitored with this sensor in yeast cells shows several distinct features: i) a variable-length of the low-FRET-state, but a similar half time of the transition to the high-FRET state, ii) the transition from the low to high-FRET aggregate state is spatially and temporally resolved akin a nucleation event, iii) later foci appearance and absence of high-FRET foci in healthier faster-dividing cells, and iv) in the later stages of aggregation, widespread condensation/aggregation occurs throughout the cytoplasm. Strikingly, while the low FRET state is more prominent in yeast than in HEK293t cells, the kinetics of the transition from the low-FRET to the high-FRET state is similar.

### DNAJB6 oligomerization behavior in the presence of mHttex1 aggregation

One of the method’s advantages is that it allows easy combination with a second genetic intervention. The chaperone DNAJB6b is an excellent case study because it prevents polyQ aggregation by inhibiting early oligomer formation.^23^ We co-transfected and expressed DNAJB6b and Q71-VC in HEK293t and compared the resulting confocal fluorescence images with Q71-VC alone (Fig 3A). Coexpression of DNAJB6b strongly reduced the number of Q71-VC aggregates, and only a few smaller aggregates remained. The FRET/donor of Q71-VC aggregates with and without DNAJB6 is similar (Supp Fig 7B), and foci with lower FRET/donor were observed with the same frequency, that is, 3% below 0.6 and 9% below 0.8, for both Q71-VC without DNAJB6 (n=117 foci) and with DNAJB6 (n=59 foci). These data suggest that the DNAJB6b does not prominently stabilize a low-FRET structure along the Q71-VC aggregation pathway.

**Figure 3.**
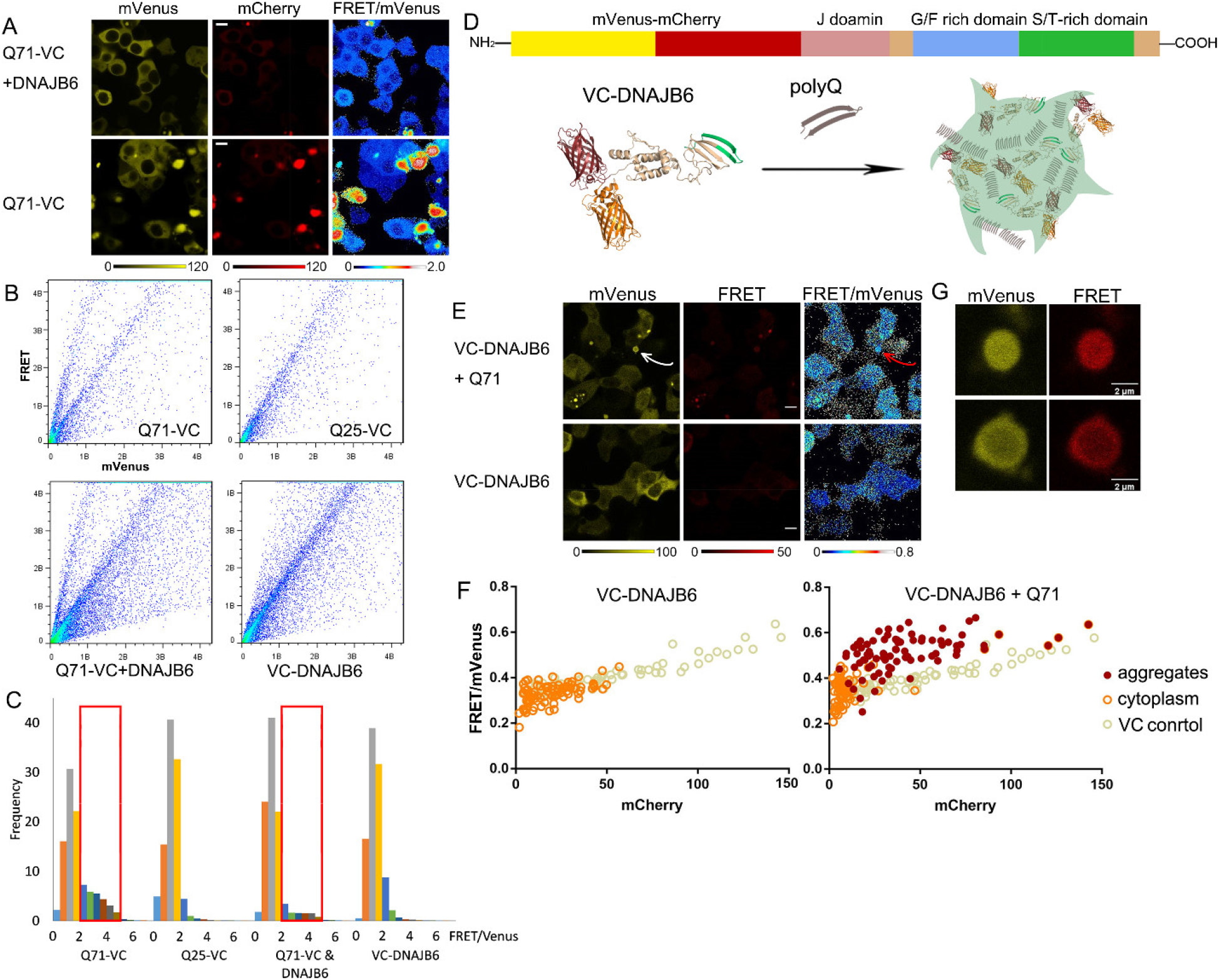
DNAJB6b suppresses mHttex1 aggregation and can form a complex with Httex1. A. Fluorescence intensities and ratiometric images of Q71-VC with and without coexpression of DNAJB6b. B. FRET versus mVenus channel of Q71-VC, Q25-VC, cotransfection with DNAJB6b, and VC-DNAJB6 in HEK293t as obtained by FACS showing clear differences in populations. C. Histograms of the individual FRET/donor ratios obtained by FACS. The red boxes indicate the FRET/mVenus ∼2-5. D. The VC-DNAJB6 design and schematic of VC-DNAJB6 inclusions in mHttex1 aggregates. E. VC-DNAJB6 co-transfected with and without Q71. Ratiometric images show an increase in FRET in the few foci. Foci and condensates are not observed without Q71 F. VC-DNAJB6 FRET/donor ratio versus the protein concentration with (right panel) and without (left panel) Q71. Q71 induces high FRET/donor aggregates. Every data point is a cell. G. Different types of aggregation complexes formed by VC-DNAJB6 and Q71. (scale bars for A and E are 10 μm.)

A powerful application of this approach is the possibility of rapid screening by fluorescence-assisted cytometry sorting (FACS) (Fig 3B). We compared the absence and presence of DNAJB6b on Q71-VC aggregation in many cells. By plotting the FRET channel versus the donor channel, we find a single main population for the Q25-VC with a slope that depends on the FRET efficiency. The Q71-VC shows a distinct additional population corresponding to the aggregates’ higher FRET/donor. We analyzed the FACS data by categorizing the FRET/donor ratios in histograms (Fig 3C). We find that the main population with lower FRET/donor is present in all experiments while the Q71-VC shows a well-populated segment with a FRET/donor >2, compared to the others. Confirming our microscopy experiments, coexpression of the DNAJB6b reduces Q71-VC aggregate formation from 29% to 11%, as assessed by the fraction with a FRET/donor higher than 2. Hence the method allows rapid and large-scale screening by FACS.

To understand the chaperone’s behavior in the absence and presence of Q71 mHttex1, we tagged the DNAJB6b with the VC to generate VC-DNAJB6 (Fig 3D). Purified DNAJB6b can form oligomers in a buffer,^24^ and tagging DNAJB6b with the VC would show how it oligomerizes in a cell and if widespread oligomerization is needed for its function. The VC was fused to the N-terminus of DNAJB6b to prevent inhibition of the active serine/threonine-rich domain located near the C-terminus of DNAJB6b.^23^ Transfection and expression lead to the full-length protein formation, as assessed by SDS-PA gel electrophoresis (Sup Fig 6). Monitoring the VC-DNAJB6b in cells by confocal microscopy gave FRET/donor ratios equal to the VC control, with a similar dependence on the concentration (Fig 3E,F). Dispersed VC-DNAJB6 is therefore indistinguishable from the monomeric VC. Further, the FRET/donor ratios were homogeneous throughout the cytoplasm, suggesting an absence of any local self-assembly.

We next studied the VC-DNAJB6 behavior with coexpression with unlabelled Q71 (Fig 3E,G,F). The Q71 did not influence the FRET efficiency of dispersed VC-DNAJB6, which remained similar to the VC. A few cells again showed foci with a similar frequency as Q71-VC with unlabelled DNAJB6b: Labelling Q71 with GFP showed that VC-DNAJB6 and Q71 colocalize in these foci (Sup Fig 6), confirming previous observations of DNAJB6 colocalization with mHttex1 aggregates.^25^ Coassembly of VC-DNAJB6 with Q71 foci is not due to VC because the control VC remains dispersed upon coexpression of Q71 (Sup Fig 6). VC-DNAJB6 in aggregates displays a higher FRET/donor of ∼0.6 than in the cytoplasm, where it has a ratio of ∼0.4. Hence, the tag provides a ratiometric readout for the coaggregation with Q71. Here, additional quenching by the Q71 fibrils could influence the ratio: The VC-DNAJB6 is locked in the aggregates, as the fluorescence did not recover after photobleaching (Sup Fig 5). Photobleaching the acceptor showed an increase in donor intensity indicative of the presence of FRET, however. The DNAJB6b distribution displays somewhat of a corona in most aggregates (∼80%), increasing with about 50% fluorescence. The FRET/donor was independent of corona formation or localization in the corona, suggesting similar environments for the DNAJB6. Together, VC-DNAJB6 can co-aggregate with Q71 aggregates, providing an increase in the FRET/donor ratio.

### FUS assemblies have mutant-dependent structures

The method distinguishes structures at ∼2-10 nm resolution: Random interacting components should not increase FRET efficiency beyond the corresponding concentration while specific interactions increase or decrease the FRET efficiency. FUS occupies mutation-dependent states in a buffer with varying RNA affinity.^6,26^ Such states become more ambiguous to categorize in cells as other molecules partition in the condensates. To this end, we measured the structural properties of foci formed by wild-type FUS and a common pathogenic mutant in living cells. In cells, wild-type FUS is transported into the nucleus, while mutations in the nuclear localization signal (NLS) retain FUS in the cytoplasm, where it forms condensates and sequesters RNA.

To assess the VC domain’s influence on FUS function, we first compared fusion at the N-terminus (VC-FUS) versus the C-terminus (FUS-VC) (Fig 4A,B). All constructs were expressed intact upon transfection in HEK293t, as shown by SDS-PAGE (Sup Fig 8). We find that fusion at the C-terminus (FUS-VC) does not localize in the nucleus and forms foci in the cytoplasm. This behavior is retained when a 13 amino acid linker is inserted between the FUS and the VC. In contrast, VC-FUS displays native-like behavior by localizing in the nucleus in a dispersed state.

**Figure 4.**
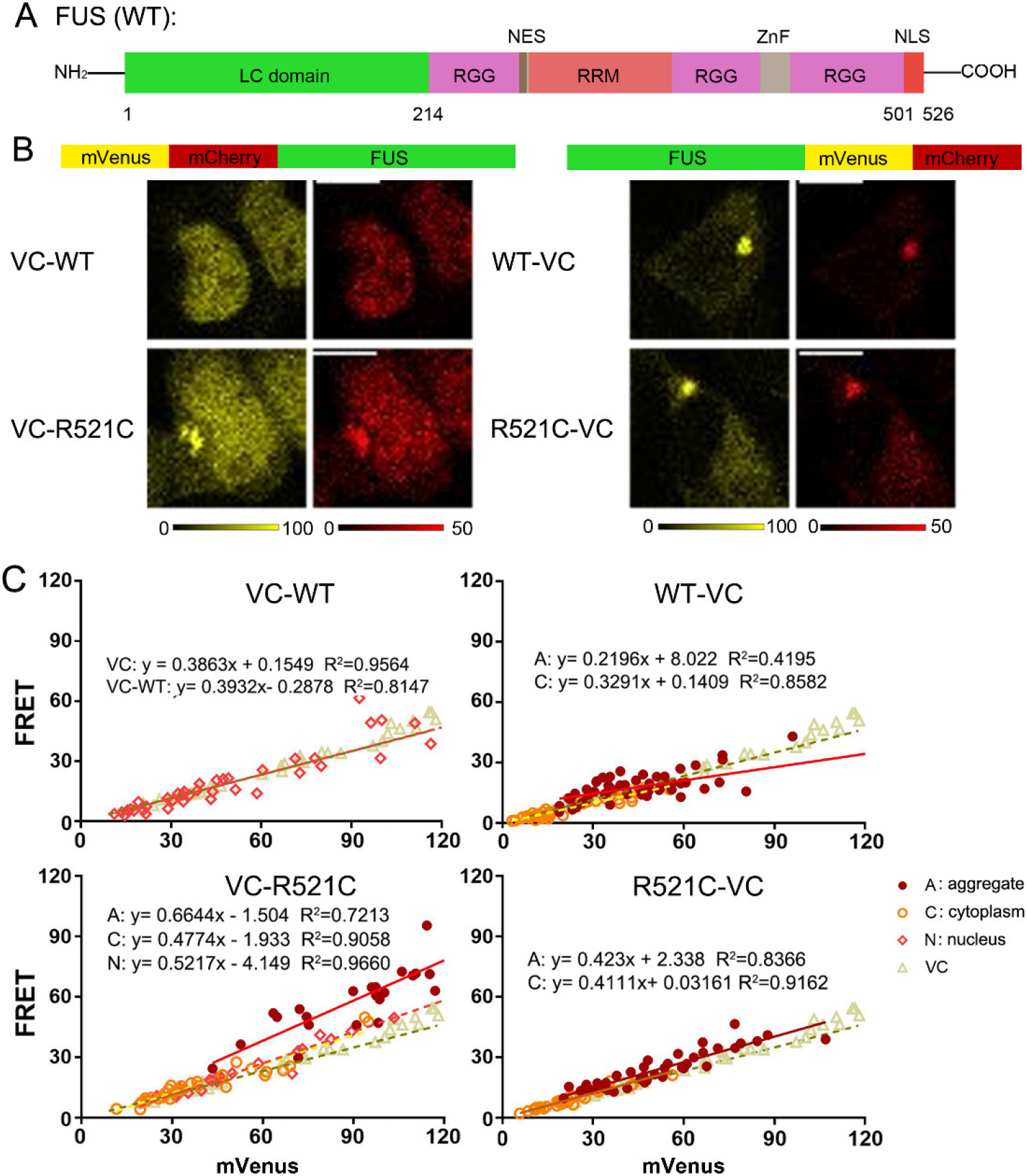
FUS mutants form foci with distinctly different structures. A. The structure of wild-type FUS. (LC: low complexity domain, NES: nuclear export signal, RRM: RNA recognition motif, NLS: nuclear localization signal) B. Probe design with VC-FUS on the left and FUS-VC on the right, and the corresponding fluorescence intensity images of HEK293t cells transfected below. Both for VC-FUS and FUS-VC, the wild-type and the R521C mutation is depicted. C. FRET plotted versus mVenus for the different FUS constructs. The data in the foci, cytoplasm, and nucleus were analyzed separately, and each fitted to a linear equation for comparison with the VC control. Each data point is a different cell. A biological replicate is displayed in Sup Fig 7. Scale bars are 10 μm.

To assess if the probing method provides information on the focus’ nanoscopic structure, we plot the FRET channel versus the donor channel: The slope of a linear fit provides the FRET/donor ratio while comparing the same concentration regimes (Fig. 4C).^27^ The dispersed VC-FUS and the control VC have a similar FRET that is linear with the concentration, showing that the VC-FUS behaves similarly to the VC alone. FUS-VC, on the other hand, formed cytoplasmic foci with a large fraction dispersed in the cytoplasm. While the FRET/donor of dispersed VC-FUS and FUS-VC overlaps with the VC control, the FUS-VC foci provide a similar or even lower FRET/donor than the VC control. Possibly, intermolecular FRET is somewhat reduced through a specific conformation or co-condensing macromolecules. Half-bleaching the FUS-VC foci lead to rapid recovery with a t1/2 of 2.2±0.8 s (s.d.,n=3) (Sup Fig 8). The foci split and fused during the experiment. Hence, the FUS-VC foci are dynamic, and the sensor shows that these condensates have mostly a random nanoscopic structure, with a slight deviation.

To obtain information on whether a common pathogenic mutation induces foci with a different nanoscopic structure, we tested the pathogenic R521C mutant associated with ALS.^28-30^ This mutant binds more tightly RNA in cells than the wild-type, leading to static complexes in a buffer,^26^ and weakens the binding with Karyopherin β2,^31^ a transportin that carries various RNA-binding proteins to the nucleus. Measurement of the VC-FUSR521C by confocal fluorescence microscopy showed that the mutation indeed induces foci formation, in contrast to the parent VC-FUS (Fig 4B). The VC-FUSR521C foci were located in the cytoplasm while a fraction remained dispersed in the nucleus and cytoplasm. Complete bleaching of the foci by FRAP lead to a rapid recovery of 2.1±1.0 s (s.d., n=2), suggesting a rapid exchange of the components with the cytoplasm. The foci were too small for accurate half-bleaching experiments. The VC-FUSR521C foci displayed a higher FRET/donor of ∼0.66 than the VC-FUSR521C dispersed in the cytoplasm or nucleus (0.48 and 0.52, respectively). The increased FRET/donor suggests that the fluorophores are situated more closely or with a favorable orientation within the focus. The R521C mutation also increases the FRET/donor in the FUS-VCR521C foci compared to the parent wild-type FUS. The FUS-VCR521C construct provided a recovery half time of 2.1±0.7 s (s.d., n=3) from FRAP by full-bleaching, suggesting a similar rapid exchange of components. Thus, FUS foci have similar partitioning dynamics but distinct mutant-dependent nanoscopic structures based on their FRET efficiencies.

## Discussion

Ubiquitous protein assembly in cell biology results in diverse and dynamic architectures that are often part of the assemblies’ biological function. Here we apply a method that allows better monitoring of the corresponding structural transitions with high spatiotemporal resolution. By fusing an mVenus-mCherry to the well-studied aggregating protein mHttex1, the chaperone DNAJB6, and the condensate-forming FUS, we show that we can distinguish between different assembly structures and can measure their interconversion resolved in space and time.

The readout is exceptionally sensitive to the assembly structure at the distance of ∼2-10 nm: The high nonlinear FRET dependence on the distance and the orientation between the fluorophores provides a unique sensitive signature of a particular structure. The relation between the FRET efficiency and structure can deviate because a fraction of ideally localized fluorophores may change the average FRET disproportionally, and competing donor quenching by fibrils may alter the readout.^13,14^ Nonetheless, a high FRET efficiency should relate to highly ordered aggregates, while unstructured condensates will have the same concentration dependence of the FRET efficiency as freely diffusing proteins. Any deviation should be the consequence of specific interactions or preferred orientations between the probed proteins, providing a fingerprint for a given assembly.

Fluorescent protein fusions influence the aggregation behavior of proteins.^32 14,15,33^ However, the VC-tagged proteins mostly recapitulate known mutation-dependent transitions and assembly behavior: i) the glutamine length dependence of the VC-Q series follows the established glutamine dependence, ii) intermediate phases in mHttex1 aggregation also have been shown to occur in a buffer without fluorescent protein fusions,^34^ iii) the VC does not dominate VC-DNAJB6 binding to Q71 because the VC control does not bind Q71, iv) the VC-DNAJB6 retains its activity by preventing Q71 aggregation, and v) the tag influences FUS condensate formation depending on the terminus tagged, while the 521C mutation always induces expected assemblies. Moreover, we use the fluorescent proteins’ monomeric versions to reduce any interactions between the fluorescent proteins themselves. Nonetheless, the fluorescent proteins’ interactions may influence protein behavior that should be assessed for each study.

With these considerations in mind, we showed that the probing method samples structures ranging from disordered condensates to fibril-type aggregates. A protein previously characterized as a condensate-forming protein, FUS, showed a low FRET efficiency similar to the VC control, as expected from a mostly random and dynamic structure. Minor mutant-dependent variations in the ratio may arise from some degree of order in the FUS assemblies, which could occur perhaps through RNA\ templating,^35^ albeit verification would require a more systematic mutagenic screen. FUS assemblies also may have some fibril content that possibly interacts with the FRET donor.^36^ Also, the mHttex1 aggregation pathway appears to embark from a more disordered assembly, especially in yeast, where a long-term low-FRET state is found. Previously, Peskett et al. showed a liquid-like state in mHttex1 assembly,^15^ which would indeed correspond to our low FRET values. Various aggregating proteins have been shown first to form a liquid-type condensate before aggregation,^37^ including mHttex1.^15^ The difference in the lifetime and the ability to resolve this state likely depends on chaperone activity and expression levels.^38^ In future studies, we aim to resolve how cell-to-cell variability in protein homeostasis underlies the correlations that we observe in yeast, between the time needed to complete one division and the kinetics with which the assemblies transition from a low-FRET to the high-FRET state. The DNAJB6-VC provides relatively low FRET/donor when coaggregating with Q71, which is relatively independent of the concentration in the corona-type structures and between aggregates. Because intermolecular FRET depends on concentration, a decreased dependence could indicate contribution by other mechanisms, such as donor quenching by fibrils. Exceptionally high FRET values are found for the mHttex1 aggregates, which have fibril-rich solid-like characteristics as indicated by SDS insolubility and FRAP experiments. Fibril-type states are structured with the fluorophores packed relatively closely, generating high FRET while providing additional donor quenching pathways. Our observation of spatially resolved aggregate (high-FRET) formation from within condensates (low-FRET) suggests a step-wise process from a condensate-like to a solid-like state.

Together, this highly sensitive method opens up an additional route to investigate the properties of condensates and aggregates in living cells. It can be exploited in a high-throughput fashion by FACS and should be amenable to provide great detail through time-resolved anisotropy measurements or fluorescence lifetime imaging. The simplicity of ratiometric imaging combined with proper controls allows straightforward access to great detail in space and time. These assets enable distinguishing structures and intermediate kinetics, benefiting the study of the assembly’s biological consequences.

## Acknowledgments

We thank Prof. Andreas Herrmann, Prof. Bert Poolman, Prof. Harm H. Kampinga, and Dr. Steven Bergink for valuable discussions. We thank Prof. Harm H. Kampinga and Dr. Steven Bergink for sharing mHttex1, DNAJB6b, and FUS plasmids. The work was funded by the ERC Consolidator Grant (PArtCell; no. 864528) to AJB, the China Scholarship Council grant to QW, the Netherlands Organization of Scientific Research (OCENW.GROOT.2019.068 en NWO-vici (VI.C.192.031)) to LMV.

